# How Does Sampling Affect the AI Prediction Accuracy of Peptides’ Physicochemical Properties?

**DOI:** 10.1101/2025.01.29.635451

**Authors:** Meiru Yan, Ankeer Abuduhebaier, Haojin Zhou, Jiaqi Wang

## Abstract

Accurate AI prediction of peptide physicochemical properties is essential for advancing peptide-based biomedicine, biotechnology, and bioengineering. However, the performance of predictive AI models is significantly affected by the representativeness of the training data, which depends on the sample size and sampling methods employed. This study addresses the challenge of determining the optimal sample size and sampling methods to enhance the predictive accuracy and generalization capacity of AI models for estimating the aggregation propensity, hydrophilicity, and isoelectric point of tetrapeptides. Four sampling methods were evaluated: Latin Hypercube Sampling (LHS), Uniform Design Sampling (UDS), Simple Random Sampling (SRS), and Probability-Proportional-to-Size Sampling (PPS), across sample sizes ranging from 100 to 20,000. A sample size of approximately 12,000 (7.5% of the total tetrapeptide dataset) marks a key threshold for stable and consistent model performance. This study provides valuable insights into the interplay between sample size, sampling strategies, and model performance, offering a foundational framework for optimizing data collection and AI model training for the prediction of peptides’ physicochemical properties, especially for prediction in the complete sequence space of longer peptides with more than four amino acids.

## 1. Introduction

Artificial intelligence (AI) has emerged as a pivotal tool for predicting the physicochemical properties of proteins and peptides, including self-assembly behaviors,^1,2^ antimicrobial properties,^3^ and stability,^4^ *etc.*, as well as properties of other materials such as high-entropy alloys,^5,6^ small molecule drugs,^7,8^ semiconductors^9^ and more. These AI models are often trained on data generated by multiscale simulations,^10–12^ in-house experiments^13–15^ or publicly available databases,^15,16^ which serve as the basis for subsequent predictions.^1,13^ However, a critical aspect that is often not addressed in such research is the overall distribution (*i.e.*, representativeness) of the training data. Specifically, in the predictions of peptides’ properties, there is a lack of justification as to whether the training data can truly represent an overall distribution of sequences or properties of peptides in the entire sequence space, namely whether the training data are biased towards certain subset sequences or properties of the sequence space. Potential biases in training data can impair the AI model’s ability to capture the intricate interplay between peptide sequences and their physicochemical properties, limiting the model’s capacity to generalize across the entire sequence space for reliable predictions.^17^

The limitation of the data distribution can be attributed to two primary factors, one being the insufficient data volume and the other being an inappropriate (less powerful) sampling approach. These two factors may act independently or together to affect the representativeness of the datasets, and subsequently the prediction accuracy. In datasets with a substantial volume of samples, the sampling method may be less critical due to the inherent diversity that the large samples tend to provide. Nevertheless, there are exceptional cases where the sampling strategy takes on increased significance. Such a case is the prediction of properties of short peptides within the entire sequence space, which boasts a staggering diversity of more than 20^50^ unique sequences (if we constrain the length of “peptide” to 50 amino acids). In such cases, it is imperative to adopt a sampling approach that ensures a sufficient and representative dataset with a minimum number of samples, capturing the distribution of both peptide sequences and their associated properties across the entire sequence space. This is essential because the exhaustive sampling required to generate a comprehensive training dataset can be prohibitively expensive in terms of computational resources or experimental costs, especially when extending the prediction to longer peptides with more than four amino acids. For instance, even pentapeptides (five amino acids) possess more than 3 million distinct sequence combinations, necessitating strategic sampling of a representative subset for AI model training and further prediction, balancing the cost of data acquisition (via computational simulations or wet-lab experiments) while ensuring reliable and rigorous predictions.

However, many studies in materials’ property prediction, including but not limited to peptides’ property-related predictions, are predicated on limited sample sizes and unjustified sampling approaches, potentially overlooking the impact of the sample volume and sampling approaches on model performance, thus the impact of sampling strategies on model accuracy remains underexplored. For instance, Batra *et al.*^1^ developed a hybrid AI system that integrates Monte Carlo Tree Search and Random Forest (RF) algorithms with coarsegrained molecular dynamics (CGMD) to efficiently predict self-assembling peptides. The AI system outperformed the traditional design of self-assembling peptides by human experts, however, the rationale behind the training data quantity and sampling strategies was not articulated, possibly resulting in high false positives in peptide self-assembly design, capping the prediction accuracy at 66.7%. In addition, Jiaqi *et al.*^18^ introduced a Transformer-based deep learning model to predict the aggregation propensity (AP) of short peptides (aggregation, a prerequisite of self-assembling). While the Transformer model provided a high-throughput approach for discovering self-assembling peptides, its performance declines with more complex systems such as decapeptides and mixed pentapeptides, compared to shorter pentapeptides. This decline can be attributed to the exponentially increasing complexity of the sequence space, necessitating a more extensive and balanced training dataset to accurately capture the distribution of peptides across the complete sequence space.

In specific cases, even with a sufficient quantity of training data, biased sampling, characterized by the overabundance of prevalent amino acid sequences and/or the deficiency of infrequent or complex peptides, may fail to capture the full diversity of peptide sequences and properties across the complete sequence space. Consequently, AI models trained on such datasets may exhibit diminished performance when extrapolating to less common or complex peptide sequences, resulting in unreliable predictions. For example, researchers frequently utilize publicly available databases ^19,20^ that comprise natural peptides and associated properties, for AI model training aimed at prediction of properties of new peptides. These natural peptides, ubiquitous in biological systems, often exhibit inherent biases towards certain amino acids. A case in point is the frequent overrepresentation of cysteine, attributable to its pivotal role in the formation of disulfide bonds, which are essential for the stabilization of peptide and protein conformations. Despite the extensive array of peptide sequences found in nature (*i.e.*, those not artificially designed and synthesized), AI models trained exclusively on natural peptide datasets are prone to inherit these biases, thereby constraining their capacity to generalize to the full diversity of the sequence space and hindering the *de novo* design of peptides for versatile applications.

In summary, the rationale behind the selection of sample size and sampling approaches in previous studies has not been sufficiently justified, potentially leading to biased training datasets and unreliable AI predictions. To address this challenge, this study aims to investigate the effect of sample size and sampling methodologies on the accuracy of AI predictions pertaining to physicochemical properties of tetrapeptides. Specifically, we explored 4 distinct sampling techniques: Latin Hypercube Sampling (LHS),^21^ Uniform Design Sampling (UDS),^22^ Simple Random Sampling (SRS), ^23^ and Probability-Proportional-to-Size Sampling (PPS).^24^ Concurrently, we have evaluated the effects of 22 sample sizes ranging from 100 to 20000 (the exact values can be found in Section 2.2 Sampling Strategies). The physicochemical properties of tetrapeptides under investigation include AP, hydrophilicity (logP), and isoelectric point (pI), which are all generated in our lab through CGMD, reported experimental data,^13–15^ and AI prediction tool,^15,16^ respectively, ensuring the consistency of the data obtained. Those three properties have significant implications in various fields such as drug design, materials science, and biotechnology, and accurate prediction of these properties through advanced computational techniques and experimental validation is essential for optimizing peptide performance and developing innovative peptide-based applications.

## 2. Methods

This study presents a three-step workflow, as illustrated in Figure 1, to evaluate the effects of sample size and sampling strategies on the AI prediction accuracy of physicochemical properties of peptides. The workflow consists of three following stages: 1) **First stage** (Figure 1a): generation of training data with CGMD simulations, available experimental data, and AI tools; 2) **Second stage** (Figure 1b): sampling within the complete sequence space of tetrapeptides, employing 4 distinct sampling methods, each yielding datasets across 22 sizes (each with 10 repetitions) ranging from 100 to 20000; 3) **Third stage** (Figure 1c): training of deep learning models using Transformer framework, for prediction of physicochemical properties of tetrapeptides. The details of each stage are provided below.

**Figure 1:**
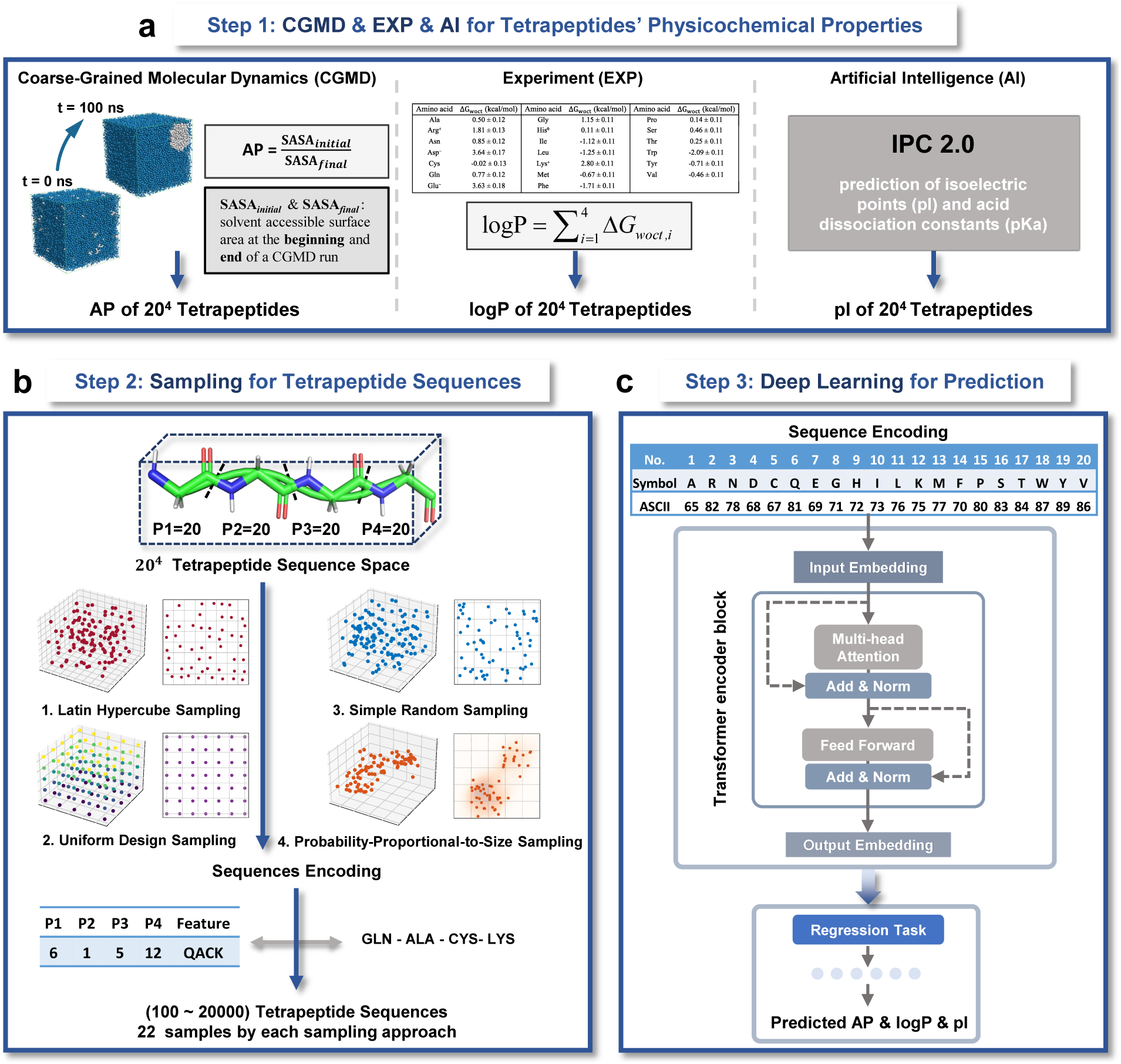
Workflow of AI model training for prediction of peptide physicochemical properties, for assessing the effects of sample size and sampling approaches on prediction accuracy. **a.** Generation of training data: aggregation propensity (AP) by coarse-grained molecular dynamics (CGMD) simulations; hydrophilicity (logP) by reported experiment data; and isoelectric point (pI) by prediction of AI models. **b.** Sampling frameworks: sampling of tetrapeptide sequences by four sampling approaches, each sampling 22 datasets with 10 repetitions, with size ranging over 100 to 20,000. **c.** Training of Transformer-based deep learning model: Prediction of physicochemical properties with Transformer-based deep-learning models, with tetrapeptide sequences encoded using ASCII code.

### 2.1 Generation of Training Data

In this study, we investigated three physicochemical properties of 160,000 tetrapeptides, namely AP, logP, and pI (Figure 1a), with the goal of deriving generalized insights into the effects of sample size and sampling method on the prediction accuracy of properties with distinct distributions (*e.g.*, normal distribution or multimodal distribution). The calculated property data are listed in the **Supplementary Data file**.

#### 2.1.1 Aggregation Propensity (AP)

To obtain the AP values of tetrapeptides, we conducted CGMD simulations with Martini force field version 2.1,^25,26^ which has been rigorously validated for its effectiveness and accuracy in assessing the aggregation of short peptides.^27^ This force field simplifies molecular representations while retaining critical interactions by approximating four atoms into a single “bead”, rendering it particularly suitable for high-throughput peptide simulations. We performed a total of 160,000 CGMD simulations in an aqueous environment, each containing 120 coarse-grained tetrapeptides of the same type and approximately 4000 water beads, equivalently to 16,000 water molecules due to the coarse-graining effect. The simulation box dimensions were set to 8 nm × 8 nm × 8 nm, yielding a peptide concentration of 0.39 mol/L and a water density of approximately 1 g/cm^3^. The simulations were conducted under *NpT* ensemble (*i.e.*, constant particle number *N*, a pressure *p* fluctuating around an equilibrium value ⟨*p*⟩ and a temperature *T* fluctuating around an equilibrium value ⟨*T* ⟩) conditions at 300 K temperature and 1 atm pressure, with each simulation running for 100 ns.

AP is defined as the ratio of the solvent accessible surface area (SASA) at the initial state (SASA*_initial_*) to that at the final state (SASA*_final_*) during a CGMD simulation,^28^ as defined in Eq. 1.

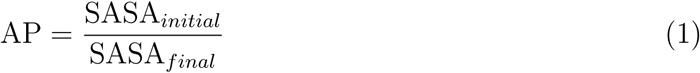

For aggregating peptides, the SASA*_final_* decreases as the simulation progresses, resulting in an AP greater than 1, while for non-aggregating, the AP values remain approximately at 1.

#### 2.1.2 Hydrophilicity (logP)

The logP data of 160,000 tetrapeptides are derived from experimental measurements,^29^ which is determined as the sum of the free energy changes associated with transferring each residue from an aqueous environment to an organic solvent of *n*-octanol. Mathematically, this relationship can be expressed as the sum of the Wimley-White whole-residue hydrophilicity (Δ*G_woct_*) as Eq. 2:

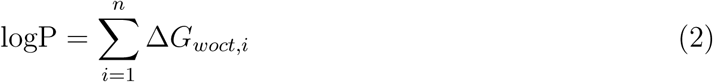

“The values of Δ*G_woct_* are calculated from the data in reference^30^ by adding the solvation energy Δ*G_glycyl_* of the -CH_2_-CONH-unit (-1.15 ± 0.11 kcal/mol^30^) to the occlusion-corrected side-chain solvation energies Δ*G*^cor^ found in Table 2 of reference^30^”. The signs of Δ*G_woct,i_*have been reversed (*i.e.*, indicating hydrophilicity) relative to those of the original publications to reflect free energies of transfer from the water phase. Here, *n* is the total number of residues in a peptide sequence (*e.g.*, *n* = 4 for tetrapeptide).

#### 2.1.3 Isoelectric Points (pI)

The pI values of 160,000 tetrapeptides were predicted with the Isoelectric Point Calculator (IPC) 2.0,^31^ a web server designed for pI and pK*_a_* (*i.e.*, acid dissociation constant) predictions based on peptide or protein sequences. Leveraging a combination of machine learning techniques, IPC 2.0 employs a separable convolution model for peptide pI prediction. The prediction models utilize ensembles of low-level models integrated with a support vector regressor, enabling accurate predictions based solely on sequence features. Compared to IPC 1.0,^32^ IPC 2.0 significantly enhances the accuracy of pI predictions.

The IPC 2.0 web server, accessible at https://ipc2.mimuw.edu.pl/, enables the highthroughput submission of peptide or protein sequences in FASTA format (a text-based format used for representing nucleotide, peptide and protein sequences).^33^ In this study, we system-atically uploaded 160,000 tetrapeptide sequences to the platform and the pI results are then predicted and output in Comma Separated Values (CSV) format for downstream processing and analyses.

The resulting CSV file contains pI values derived from 21 publicly available methods of pI prediction. In the study by Lukasz *et al.*,^31^ it was found that the Patrickios’s method exhibited the maximum observed outlier count (22,818 out of a total of 29,774 sequences) among all methods evaluated on the peptide dataset with R^2^ value less than zero, making it the least performing method among all, as shown in Table 2 of reference.^31^ To avoid introducing distinct variations and unreliability, we excluded the data from the Patrickios’s method. Subsequently, we implemented a data processing procedure to remove the maximum and minimum pI values of each tetrapeptide and obtained trimmed mean pI values averaged over the remaining 18 databases (21-1*_Patrickios_*-1*_max_*-1*_min_*) predicted by IPC 2.0.

### 2.2 Sampling Strategies

This study employs four sampling techniques: 1) LHS,^21^ 2) UDS,^22^ 3) SRS,^23^ and 4) PPS,^24^ to conduct a comprehensive analysis of how these sampling strategies influence the distribution of training data and subsequently affect the accuracy of AI predictions. Before sampling, each of the 20 amino acids is encoded with a unique numerical representation from 1 to 20 (Figure 1c). The sampling process generates tetrapeptide sequences across 22 sample sizes, *e.g.*, 100, 200, 300, 400, 500, 700, 900, 1100, 1500, 2000, 2500, 3000, 4000, 5000, 6000, 8000, 10000, 12000, 14000, 16000, 18000, 20000, with each size undergoing 10 iterations to ensure robustness and reliability. We allow for sampling with replacement, meaning that the same amino acid can appear multiple times within one tetrapeptide sequence. The sampling is performed with our in-house MATLAB and R studio codes, reposited as the **Supplementary Code file**.

It should be noted that during sampling procedures, to simplify peptide notations, we use 4-digit ABCD (*i.e.*, random letters here) to represent the peptide NH^+^-ABCD-COO*^−^*, *i.e.*, the peptide ABCD is connected with -NH^+^ group at the left end (A amino acid), and is connected with the -COO*^−^* group at the right end (D amino acid), thus the peptide ABCD is different from the peptide DCBA (*i.e.*, NH^+^-DCBA-COO*^−^*), due to the existence of different chemical groups connected at the two ends.

#### 2.2.1 Latin Hypercube Sampling (LHS)

LHS is a widely used stratified sampling (or space-filling design) method that was introduced by Michael McKay of Los Alamos National Laboratory in 1979.^34^ This method ensures that each dimension of the sample space is divided into equal-probability intervals, thereby promoting a more uniform coverage compared to SRS. This stratification approach improves the representativeness of the sample, particularly in high-dimensional spaces. In this study, LHS is specifically employed to achieve comprehensive coverage of the tetrapeptide sequence space, which consists of 160,000 combinations of 20 standard amino acids.

#### 2.2.2 Uniform Design Sampling (UDS)

Borrowing an algorithm from number theory to generate a uniform sequence, UDS is a deterministic space-filling design method that ensures sample points are distributed as uniformly as possible across the sample space. ^22,35^ Compared to orthogonal design, UDS loosens the requirement on the orthogonality and thus requiring fewer points.^36^ This method was first proposed by Kaitai Fang and Yuan Wang^37,38^ for experiments where the cost of running a trial is too high. UDS is commonly used in computer simulations and other experimental designs where uniformity is crucial for adequately covering the entire parameter space. In this research, the UDS is employed to ensure that the percentage of each amino acid at each position remains at 5%, regardless of the sample size.

#### 2.2.3 Simple Random Sampling (SRS)

SRS is a basic and straightforward sampling approach in survey sampling, in which each amino acid at each position has an equal and independent probability (*i.e.*, = 5% = 1*/*20) of being selected. While SRS is an unbiased sampling method, its intrinsic variability may lead to inadequate representation of the population probability of each amino acid, particularly when sample sizes are small. This limitation can result in unrepresentative samples, which can be detrimental to the prediction accuracy of AI models.

#### 2.2.4 Probability-Proportional-to-Size Sampling (PPS)

PPS is a probabilistic method in which samples are randomly drawn based on their assigned size. The higher the size assigned to an element, the greater its probability of being selected. In this research, each amino acid at each position is assigned a random size, and thus the PPS is essentially a SRS sampling.

### 2.3 AI Model Construction

#### 2.3.1 Sequence Encoding

The reliable encoding of peptide sequences is essential for effective model training and accurate predictions. This study employs ASCII encoding (Figure 1c) to provide additional insights into the impact of the encoding methods on prediction accuracy of peptide properties, thereby complementing the one-hot, sequence, and graph encodings that have been previously explored.^39,40^ ASCII encoding translates peptide sequences into numerical representations, which facilitates efficient processing by computational models. Specifically, each amino acid is represented by a 3-digit code corresponding to its three-letter representation. For example, the amino acid Phe is represented by the numerical sequence “80 72 69”, and each amino acid in a sequence is connected by the hyphen “-”, corresponding to the ASCII code of “45”. As a result, each tetrapeptide is encoded by a sequence of 15-bit ASCII codes (*i.e.*, 12 bits of amino acids plus 3 bits of hyphens).

#### 2.3.2 AI Model Training and Evaluation

This study employs transformer-based architectures,^41^ optimized for sequence-based predictions. Recent research has demonstrated the effectiveness of Transformer architecture in predicting peptide properties.^42–45^ The algorithm of the deep learning model training and prediction process is depicted in Figure 1c, showcasing the sequence encoding and regression tasks.

The prediction tasks focus on three properties, *i.e.*, AP data derived from CGMD simulations, experimentally obtained logP data, and AI-predicted pI data. Samples of varying sizes - 1500, 2000, 5000, 8000, 12000, 16000, and 20000 - are generated under four different sampling methods. Each sample size undergoes ten rounds of repeated sampling to assess the variability. This process results in a total of 280 datasets (7 sample sizes × 4 sampling methods × 10 iterations) for training and prediction.

Each dataset is partitioned into training, validation, and testing subsets, using a structured splitting strategy. The training and validation datasets account for 80% of the total dataset, while the testing subset comprises the remaining 20%. Within the 80% allocated to training and validation, 80% is used for training (equivalent to 80% × 80% = 64% of the total dataset), and the remaining 20% is allocated for validation (equivalent to 80% × 20% = 16% of the total dataset).

The performance of the transformer-based model was assessed on the test data set using three commonly employed regression metrics: Mean Absolute Error (MAE), Mean Squared Error (MSE), and R-squared (R^2^), which are averaged results over ten rounds of repeated sampling. MAE is used to assess the predictive accuracy of models on continuous data by measuring the average absolute difference between the predicted and actual values. MSE measures the average of the squared differences between predicted and actual values, with smaller MSE values indicating better accuracy of the model prediction. R^2^ represents the proportion of variance in the dependent variable that is explained by the independent variables in the model. R^2^ values closer to 1 indicate better model fit, as they suggest that the model explains a higher proportion of the variability in the target variable.

## 3 Results

### 3.1 Sequence Distributions

To evaluate the distribution of amino acids within tetrapeptide sequences, we analyzed the percentage distribution of four amino acids at each position (*i.e.*, Position 1 - P1, Position 2 - P2, Position 3 - P3, Position 4 - P4) across varying sample sizes (*i.e.*, *N* = 100, 500, 2000, 5000, 8000, 10000, 16000, and 20000), generated using the four aforementioned sampling approaches. It is important to note that, although each sample size with each sampling approach was repeated ten times, the results presented in this section are based on one representative repetition. The percentage distribution of each amino acid at each position is defined in Eq. 3:

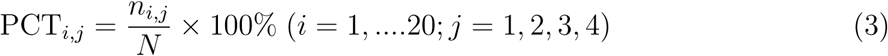

Here, PCT*_i,j_*denotes the percentage of the *i* -th amino acid at the *j* -th position, n*_i,j_*represents the total count of *i* -th amino acid occurring at the *j* -th position, and *N* indicates the total number of samples.

The heatmaps (Figure 2a-d) provide an intuitive illustration of the percentage difference in sequence distributions across various sample sizes and sampling approaches. For all methods except UDS, the percentage differences gradually decrease as the sample size increases. In contrast, the UDS method consistently maintains identical percentages for each amino acid at each position(= 5%), irrespective of the sample size. While these observations align with expectations, it is noteworthy to examine the specific sample sizes or sampling approaches (excluding UDS) at which the differences of the maximum and minimum percentage converges to zero or a specific threshold (*e.g.*, 1%).

**Figure 2:**
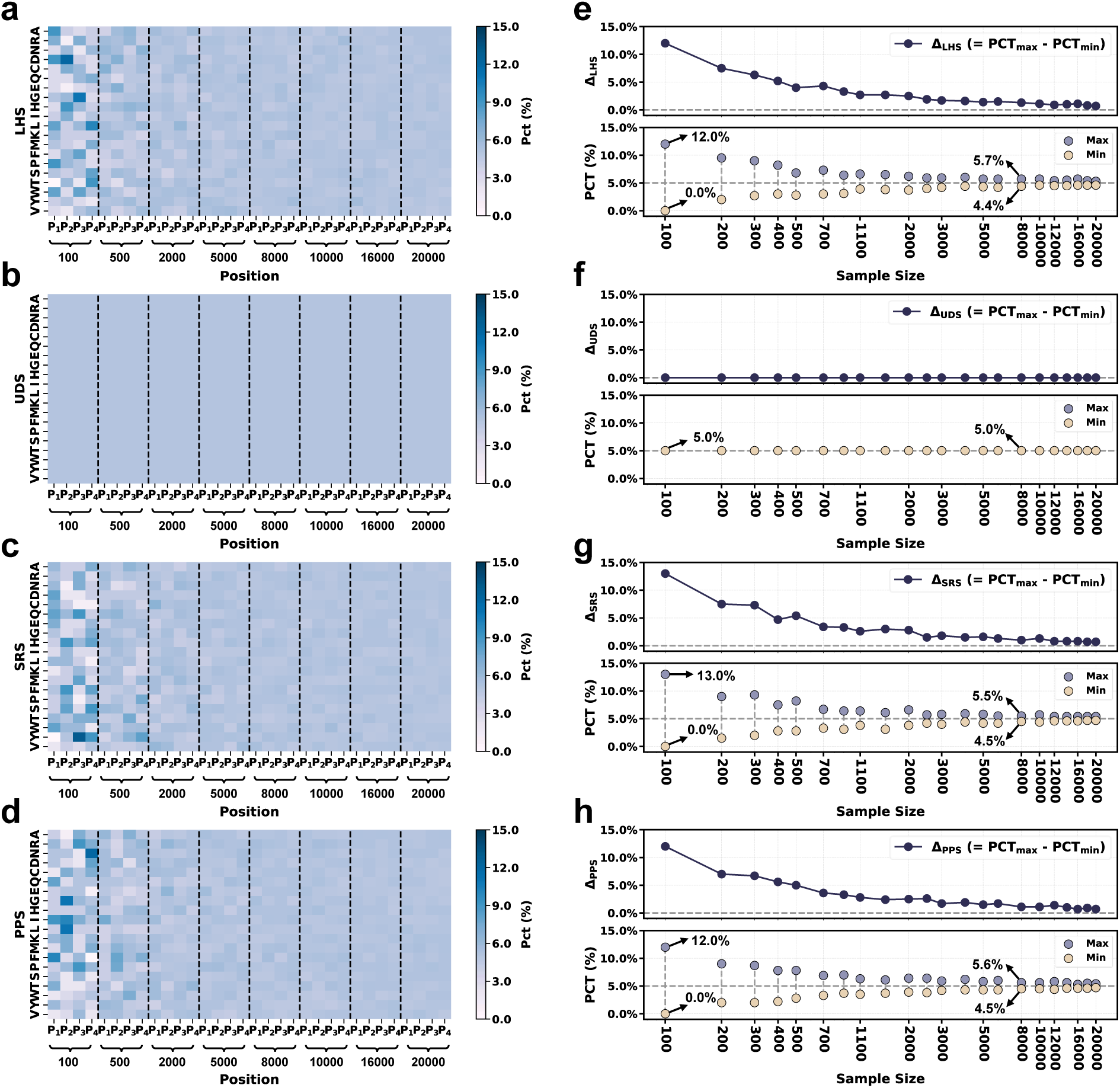
**a-d.** Percentage (PCT) distribution of 20 amino acids at four positions of P1, P2, P3, and P4, within sample sizes of *N* = 100, 500, 2000, 5000, 8000, 10000, 16000, and 20000, achieved with four sampling approaches of **a.** Latin Hypercube Sampling (LHS), **b.** Uniform Design Sampling (UDS), **c.** Simple Random Sampling (SRS), and **d.** Probability-Proportional-to-Size Sampling (PPS). **e-h.** Maximum and minimum percentage of amino acid (regardless of position), as well as the discrepancy between the maximum and minimum percentage (Δ = PCT_max_ − PCT_min_) within each sample size, of sampling approaches of **e.** LHS, **f.** UDS, **g.** SRS, and **h.** PPS. In summary, the UDS sampling consistently maintains a uniform distribution of amino acids across all sample sizes, while LHS and SRS exhibit greater variability at smaller sample sizes.

When the sample size is as small as 100, certain amino acids are not represented at specific positions in sequences, which can be generally observed with sampling methods of LHS, SRS, and PPS, resulting in 0% occurrence for these amino acids. Additionally, the maximum percentage of specific amino acids can reach 13% with the SRS method and 12% with LHS and PPS, leading to percentage difference of 13% and 12%, respectively. In contrast, the percentage difference of UDS remains at 0% (*i.e.*, no maximum and minimum can be detected), highlighting its superiority in generating more representative samples with small sample sizes. As the sample size increases to 8000, the maximum percentage difference decreases to 1.3% for LHS, 1% for SRS, and 1.1% for PPS, demonstrating reduced variability in amino acid distribution as the sample volume grows. Notably, there is no discernible difference among these three sampling methods at larger sample sizes of 8000 and more.

In summary, the UDS method is recommended for sampling smaller datasets, as it ensures equitable representation of all amino acids and thus eliminate potential biases in sequence distributions. Increasing the sample size to 8000 - equivalent to 5% of the entire sequence space of tetrapeptide - can also effectively mitigate biases introduced by other sampling methods, such as LHS, SRS, and PPS. This 5% threshold was obtained in terms of sequence space of tetrapeptide, whether it can be generalized to sequence space of longer peptides remains uncertain. It is highly likely that a higher threshold would be required to achieve a diverse distribution of training sequences, due to the exponentially increased sequence diversity. For longer peptides such as pentapeptide, even 5% of sequence space comprises 160,000 sequences, and increasing the sample size to such an extent to achieve an acceptable distribution (*e.g.*, percentage difference among amino acids and each positions smaller than 1%) would result in significantly higher costs in terms of human labor, materials, and time. These factors must be carefully considered when designing experiments. Given these challenges, UDS emerges as the optimal sampling approach for balancing representation diversity and resource efficiency.

Nonetheless, the question of whether representative sampling of sequences within training datasets guarantees unbiased sampling of their physicochemical properties still remains unresolved. To address this question, we systematically quantify the discrepancies in the distribution of physicochemical properties associated with sampled sequences across varying sample sizes and sampling methodologies in Section 3.2.

### 3.2 Property Distributions

To quantify the distribution discrepancy of the properties between the sampled sequences and the total sequence space, the errors in the property distribution, denoted by Δ, are calculated using a **three-step** process (see details of the “Calculation Steps of Average Errors of Property Distribution” in Supplementary Materials - 1). To ensure reliability, we totally perform 10 repetitions of sampling for each sample size with each sampling method, and we obtain the average of errors, standard deviations, the maximum and the minimum of the errors of the 10 repetitions.

The error of each repetition is denoted by Δ*_R_*, while the average is denoted by Δ with subscript of specific property and superscript of sampling method, for example, Δ^LHS^, or denoted simply by Δ*_mean_*. The details of Δ*_R_* with respect to each bin, repetition, sample size, sampling approach, and property are summarized in Supplementary Materials - 2. Meanwhile, the mean and standard deviation, the maximum and the minimum of 10 Δ*_R_*’s are shown in Figure 3 (bin size *k* = 10) and Figure S1 (bin size *k* = 20) in Supplementary Materials - 1.

**Figure 3:**
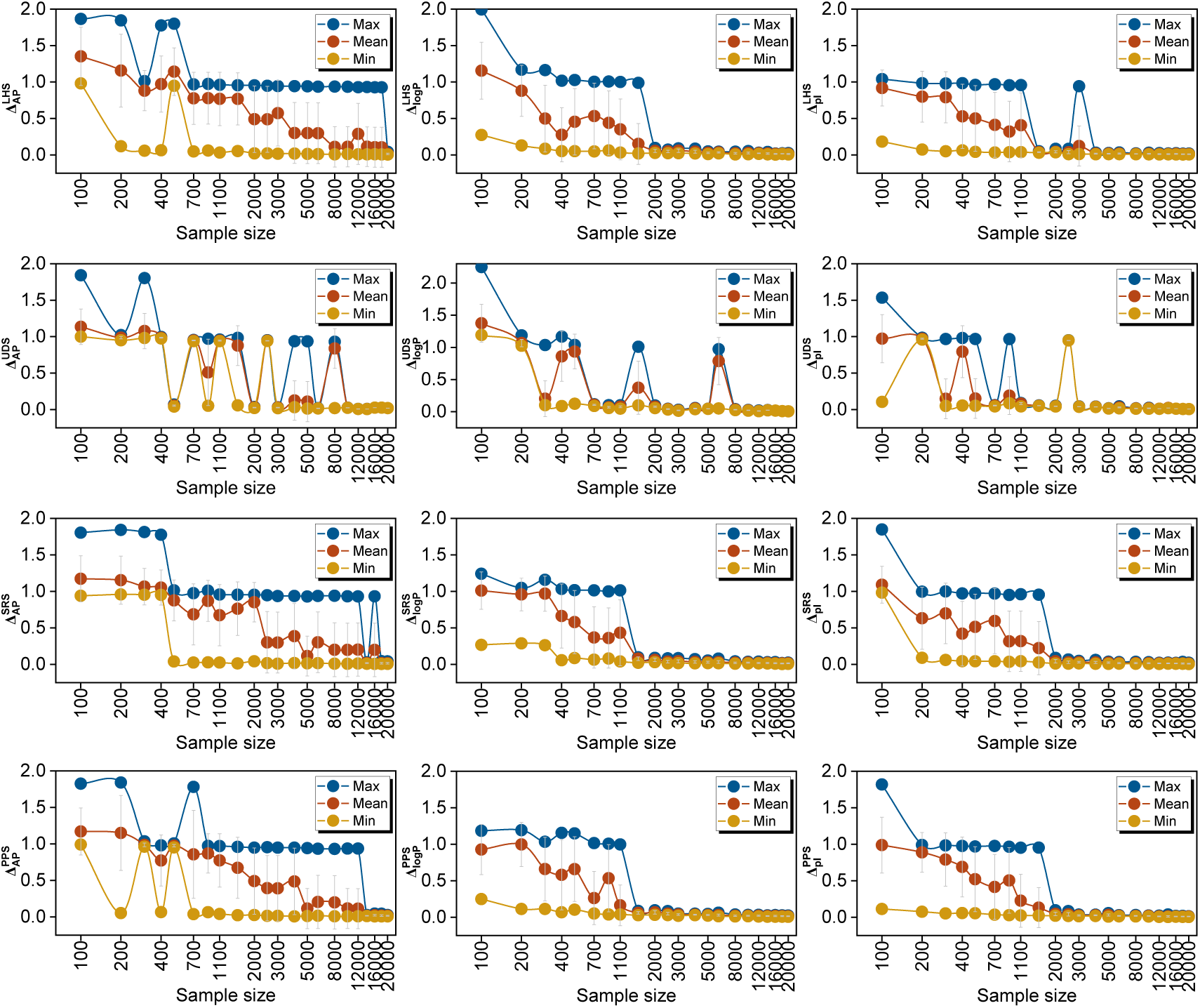
Absolute average property errors (Δ values) between the samples and the entire sequence space with bin size ***k* = 10**, including mean values of 10 repetitions (in red), maximum values (in blue) and minimum values (in yellow) among the 10 repetitions, with respect to each property, sample size, and sampling approach. The error bars in grey represent the standard deviations.

It is generally observed that the errors in the properties are predominantly located at the lower and higher ends of the distribution, particularly the higher ends (Supplementary Materials - 2), especially when the sample size is less than 8000. These errors are the primary contributors to the discrepancies depicted in Figure 3 and Figure S1 (Supplementary Materials - 1). Such errors arise due to the absence of peptide sequences with properties within specific property range incorporated in the complete sequence space, even with UDS method.

Below, we analyze the performance of each sampling method in representing the diversity of each property with increasing sample size, from two perspectives, one is the minimum sample size with an acceptable error (Δ*_mean_ <* 0.15), and the other is the minimum sample size after which the error remains stable with no abrupt increase. The criteria of “abrupt increase” is 10% increase compared to the errors of the minimum sample size with Δ*_mean_ <* 0.15. The sample sizes with abrupt increases (denoted by “10% ↑”) are also illustrated in Table 1.

**Table 1:**
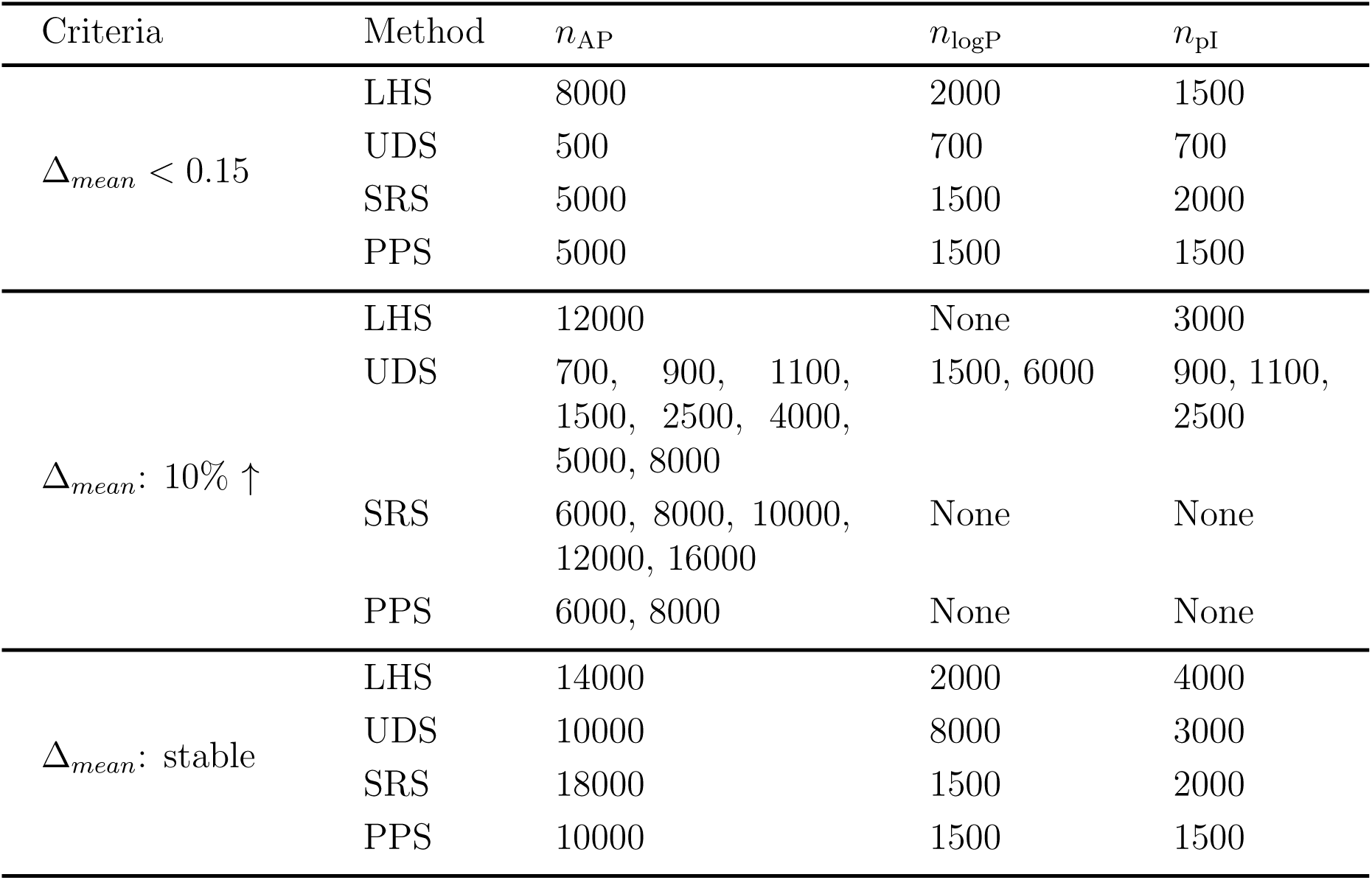
Smallest ample sizes with mean errors smaller than 0.15, sample sizes exhibit over 10% increase of error compared with that of the samllest sample sizes with mean errors smaller than 0.15, as well as sample sizes achieving stable error performance.

| Criteria | Method | $n_{AP}$ | $n_{\log P}$ | $n_{pI}$ |
| --- | --- | --- | --- | --- |
| $\Delta_{mean} < 0.15$ | LHS | 8000 | 2000 | 1500 |
|  | UDS | 500 | 700 | 700 |
|  | SRS | 5000 | 1500 | 2000 |
|  | PPS | 5000 | 1500 | 1500 |
| $\Delta_{mean}: 10\% \uparrow$ | LHS | 12000 | None | 3000 |
|  | UDS | 700, 900, 1100, 1500, 2500, 4000, 5000, 8000 | 1500, 6000 | 900, 1100, 2500 |
|  | SRS | 6000, 8000, 10000, 12000, 16000 | None | None |
|  | PPS | 6000, 8000 | None | None |
| $\Delta_{mean}: \text{stable}$ | LHS | 14000 | 2000 | 4000 |
|  | UDS | 10000 | 8000 | 3000 |
|  | SRS | 18000 | 1500 | 2000 |
|  | PPS | 10000 | 1500 | 1500 |

**Table 2:**
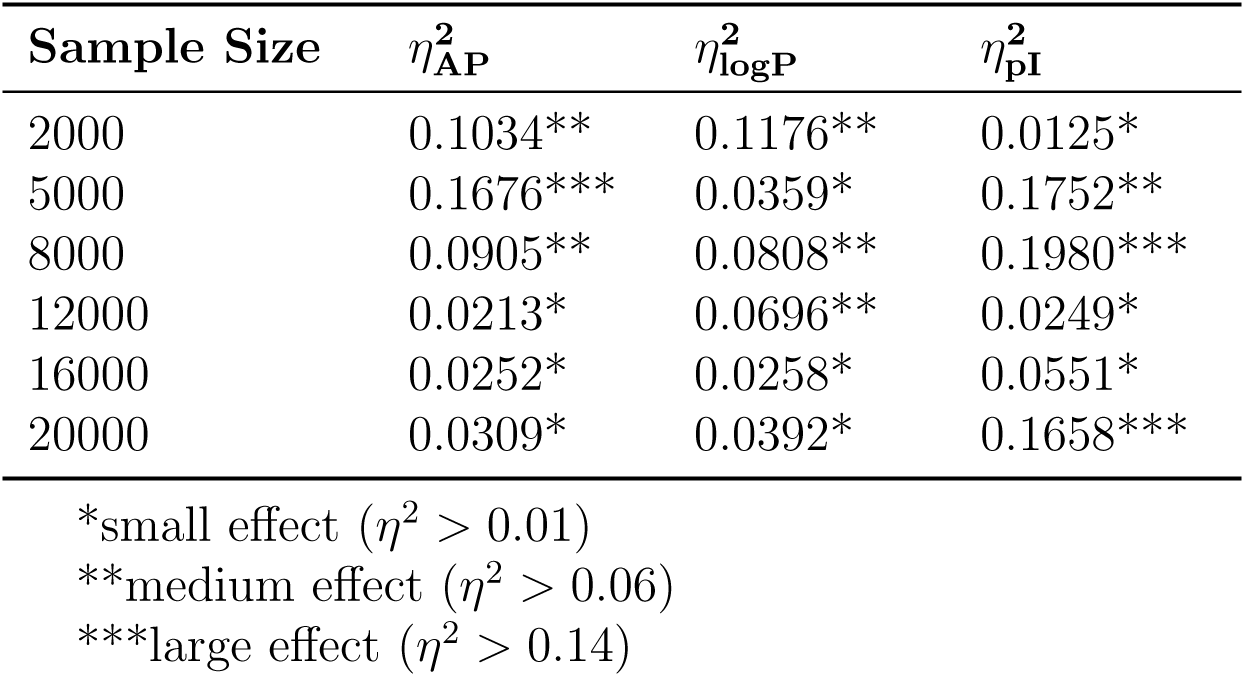
Effect size analysis (*η*^2^) across various sample sizes for AP, logP, and pI.

For the property AP, it is interesting to note that the LHS method exhibits a “ladder” decrease after the sample size of 500 (700 ∼ 1500: 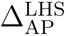 ≈ 0.7725; 2000 ∼ 3000: 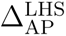 ≈ 0.5172; 4000 ∼ 6000: 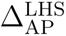 ≈ 0.2987, 8000 ∼ 20000 excluding the size of 12000: 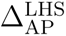 ≈ 0.0931). The 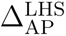 reaches 0.1102 (*<* 0.15) at the sample size of 8000, while an abrupt increase to 0.2886 appears at the size of 12000, and finally it reaches stability at the sample size of 14000. The UDS method achieves a mean error that is close to 0 (*i.e.*, 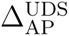 = 0.054) at the sample size of 500, with a standard deviation that is also close to 0 (*i.e.*, 0.008). However, it would be premature to conclude that the UDS method is superior in sampling properties with small errors, as this could merely be a “lucky” event - the UDS method coincidentally samples sequences with AP values across all ranges at a sample size of 500. As the sample size increases, the performance of the UDS method exhibits instability. For instance, at sample sizes of 700, 900, 1100, 1500, 2500, 4000, 5000 and 8000, the 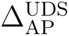 increased over 10%. This can be attributed to the fact that the properties associated with those sampled sequences do not cover the entire range, particularly the higher end of the distribution (AP*^′^ >* 0.8). Starting from the sample size of 10000, 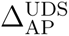 remains stable at approximately or below 0.025, indicating that the sampled sequences can cover the properties across all ranges. Therefore, it is reasonable to conclude that although the UDS method may be capable of sampling diverse properties with a small sample size, it inherently possesses instability regarding errors due to its deterministic nature - the samples are identical at each repetition, and the properties in each range may not be sampled, thus, the errors cannot be diminished by averaging the Δ*_R_* across all the repetitions. For SRS and PPS methods, the errors decrease significantly to below 0.15 with size of 5000, while even with size 16000 and 8000 for SRS and PPS, there is still a chance of bearing large errors, and 18000 and 10000 is determined as “stable” sample size, respectively.

For the property logP, the sampling error generally decreases to below 0.15 once the sample size reaches 2,000. Notably, the error drops to below 0.15 even at a sample size of 700 when using the UDS method. A similar trend is observed for the isoelectric point (pI), where sampling errors are relatively large at smaller sample sizes but become less significant as the sample size increases to 2,000. However, it is important to note that both LHS and UDS exhibit instability at larger sample sizes. For example, instability is observed at a sample size of 3,000 for pI using the LHS method, and at sample sizes of 1,500 and 6,000 for logP using the UDS method. Additionally, instability is noted at sample sizes of 900, 1,100, and 2,500 for pI using the UDS method.

Achieving a generalized conclusion on the optimal sample size to minimize errors is challenging, as it is highly dependent on both the specific property being examined and the sampling method employed. For AP, a sample size of 10,000 (6.25% of the total sequences) appears to be optimal for achieving stable errors using UDS and PPS. However, larger sample sizes of 14,000 and 18,000 are required for LHS and SRS, respectively. For logP and pI, a sample size of 2,000 is sufficient to achieve stable errors using SRS and PPS. For logP, the sample size needs to be increased to 8,000 when using UDS, while for pI, a sample size of 3,000 is required when using LHS. These observations suggest that the reliability of the property data significantly influences the sample size needed to achieve stable errors. Importantly, it is crucial to examine the AI prediction accuracy for these properties across various sample sizes and sampling methods. This analysis is presented in Section 3.3, where the impact of different sampling strategies on AI model performance is further explored.

### 3.3 AI Prediction Accuracy

Figure 4 presents the predictive performance of Transformer-based AI models in predicting the three physicochemical properties of AP, logP, and pI, using four sampling strategies with varying sample sizes ranging from 100 to 20000. Detailed results for all sample sizes are provided in Tables S1 to S4 in the Supplementary Materials - 1. The performance metrics for specific sample sizes (2,000, 5,000, 8,000, 12,000, 16,000, and 20,000) are illustrated in Figure 4.

**Figure 4:**
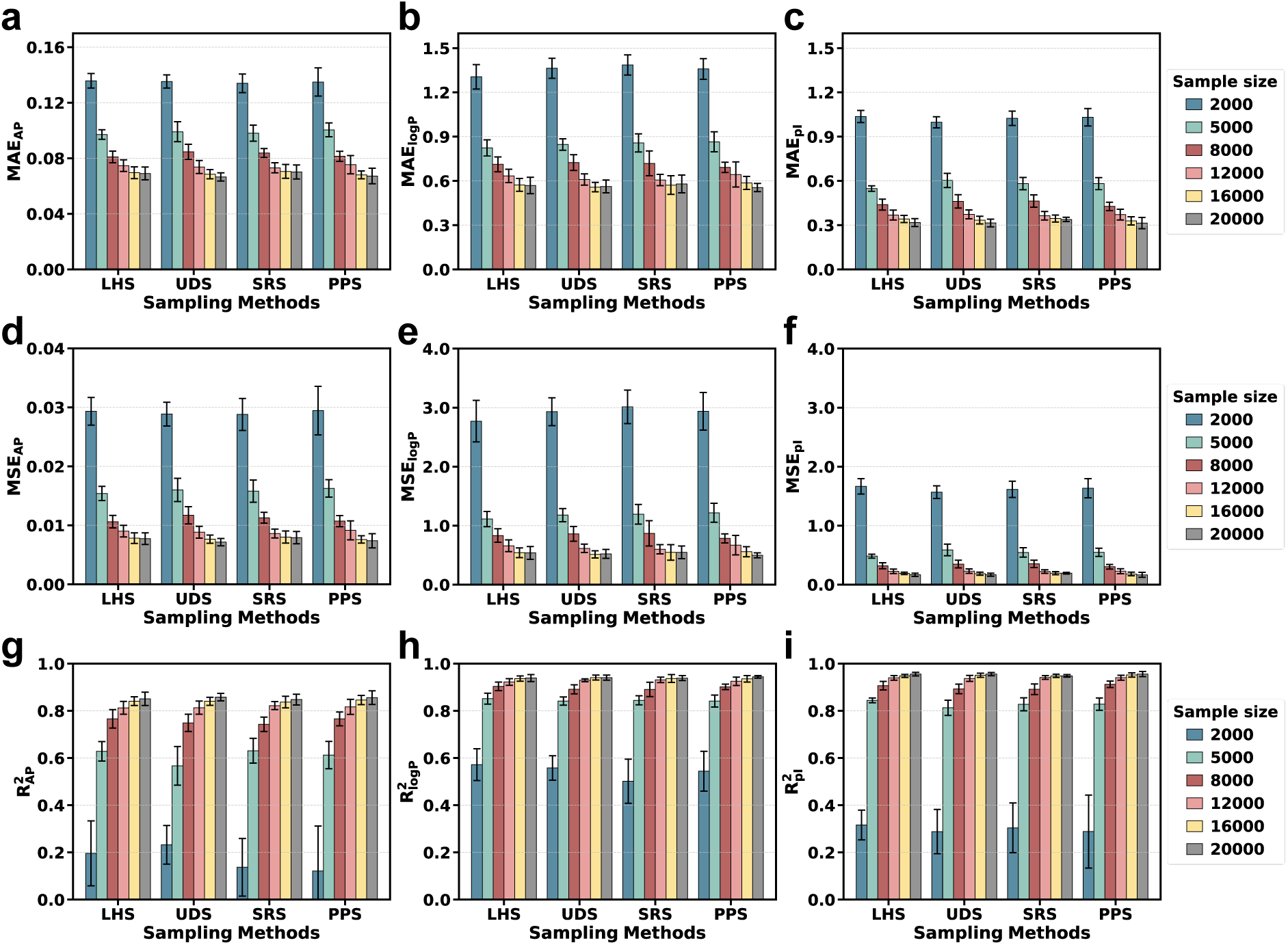
Performance comparison of four sampling methods of LHS, UDS, SRS, PPS across three physicochemical properties: AP, logP, and pI. Panels (a - c) display MAE values, (d - f) show MSE values, and (g - i) present the R^2^ values, evaluated over sample sizes of 2000, 5000, 8000, 12000, 16000, and 18000. Error bars represent the mean and the standard deviations.

Across all sampling methods, the predictive performance for the three physicochemical properties of AP, logP, and pI generally improves with increasing sample size. A significant enhancement in performance is first observed at a sample size of 5,000, followed by more modest improvements as the sample size increases from 5,000 to 20,000. Beyond a sample size of 12,000, MAE, MSE, and R^2^ stabilize, indicating that further expansion of the dataset yields diminishing returns in predictive accuracy. The stabilization of these error metrics at a sample size of 12,000 (equivalent to 7.5% of the complete sequence space) suggests that this sample size represents an optimal balance between computational cost and predictive accuracy. Additionally, it is observed that prediction errors are predominantly located at the extreme ends of the property distributions (Figure S2, Supplementary Materials - 1). This phenomenon can be attributed to the fact that sequences with properties at these extremes are often underrepresented in the sampling process, as illustrated in the error analysis provided in Supplementary Materials - 2. Therefore, potential future work could focus on incorporating more sequences with extreme property values into the sampling process. For instance, this could be achieved by assigning higher weights to peptides with extreme property values using the PPS sampling method. This approach may help improve the representation of these sequences and enhance the overall predictive performance of the models.

After verifying that the assumptions of normality and homogeneity of variances are satisfied, a One-Way Analysis of Variance (ANOVA, see details in Supplementary Materials - 1) is performed to assess the significance of the testing R^2^ of AP, logP, and pI across sample sizes of 2000, 5000, 8000, 12000, 16000, and 20000 using LHS, UDS, SRS and PPS methods. The ANOVA results indicate that no significant differences in testing R^2^ values among the groups (here “group: indicates each sampling method group with all sample sizes with respect to each property), with all F-statistics yielding p-values greater than 0.05. Although the statistical tests show no significant differences among groups, this does not necessarily imply the absence of actual differences. The lack of statistical significance could be attributed to factors such as insufficient sample size (with only 10 repeated sampling-derived R^2^ values per sample size), high within-group variability (due to sample size effect), or limitations in the chosen statistical method. Notably, even when differences are not statistically significant, variations in R^2^ values can hold biological relevance.

To further quantify the practical significance of group differences, an “effect size” analysis is conducted. Eta Squared (*η*^2^, see details in Supplementary Materials - 1) is calculated to measure the proportion of total variance explained by the group differences, even in the absence of statistical significance. As shown in the **Table 2**, the effect size provides valuable insights into the magnitude of the differences among groups, highlighting their potential biological and predictive importance.

The results reveal that sample size plays a pivotal role in determining the accuracy and stability of model predictions. At smaller sample sizes (*<* 8000), model performance is highly sensitive to the choice of sampling approach. As the sample size increases, model errors gradually converge, and the differences between sampling approaches diminish, aligning with the Central Limit Theorem principles [xx]. Specifically, a sample size of approximately 12000 marks a clear improvement in model accuracy and convergence for all three parties. Beyond this threshold, further increasing the sample size to 20000 results in diminishing returns, as the performance gains plateau. The stabilization of error metrics at 12,000 samples, equivalent to 7.5% of the tetrapeptide sequence space (20^4^), suggests a potential balance point between computational cost and predictive performance. However, it is important to note that this observation is not definitive. Further experiments on more extensive and more complex peptide spaces, such as pentapeptides and mixed peptide systems, are required to validate this hypothesis and to generalize the findings.

## 4 Discussions

This study investigates the intricate interplay between sampling approaches, sample size, and AI model performance in predicting peptides’ physicochemical properties. Despite the distinct sequence sampling strategies are employed, no significant differences in AI prediction accuracy are observed among all four sampling methods. However, from the authors’ perspective, the UDS method is still recommended to ensure sequence representativeness. Regarding the representativeness of the associated properties of sampled sequences, the UDS method may not fully account for such considerations. In contrast, the PPS method may incorporate these factors by assigning higher weights to peptide sequences with extreme values.

Rather than the sampling method, sample size emerges as a critical determinant of prediction accuracy. A sample size of approximately 8000 marks a threshold for significant performance improvement, while 12,000 samples appear to represent a practical balance point between computational efficiency and accuracy. This finding suggests that a sample size of 12000, which corresponds to 7.5% of the complete sequence space of tetrapeptides, may be sufficient for reliable predictions

These findings hold significant implications for the design of AI-driven peptide discovery pipelines, particularly in the context of drug development and biomaterial design. By optimizing sampling strategies, researchers can achieve high predictive accuracy while reducing computational costs. Although this study focuses on tetrapeptides, future work could extend these findings to longer peptides and more complex peptide systems, such as pentapeptides and mixed peptide sequences, thereby further validating the generalizability of the proposed sampling strategies.

## 5 Conclusions

This study systematically evaluated the performance of a Transformer-based AI model in predicting three physicochemical properties of AP, logP, and pI, using four distinct sampling strategies of LHS, UDS, SRS, and PPS. The results underscore the critical role of sample size, rather than the specific sampling method, in achieving reliable and optimal predictive performance. A sample size of approximately 12,000, representing 7.5% of the total tetrapeptide dataset (160,000 sequences), emerges as a key threshold. At this level, the model exhibits stable and consistent performance, suggesting that training and prediction on at least 7.5% of the dataset may be necessary to ensure reliable results. Beyond this threshold, improvements in predictive accuracy diminish, indicating diminishing returns from further increasing the dataset size. This finding provides a practical guideline for balancing computational efficiency with predictive accuracy, particularly in large-scale bioinformatics applications.

## Acknowledgement

Dr. Jiaqi Wang acknowledges the funding support of the National Natural Science Foundation of China (No. 52101023), Fundamental Research Plan (Natural Science Foundation) - General Programme of Jiangsu Provincial Department of Science and Technology (No. BK20241816), and Research Development Fund (RDF) of Xi’an Jiaotong-Liverpool University (RDF-24-01-013). Dr. Haojin Zhou acknowledges the funding support of RDF of Xi’an Jiaotong-Liverpool University (RDF-23-01-073). The authors also acknowledge the high-performance computing platform at Xi’an Jiaotong-Liverpool University.

## Supporting Information Available

Four Supplementary Materials are available: Supplementary Materials - 1 containing additional figures, methods or discussions related to the main text; Supplementary Materials - 2 containing the errors of each bin; Supplementary Data file containing the property data of 160000 tetrapeptides; and Supplementary Code file containing the four sampling codes of LHS, UDS, SRS, and PPS.

## TOC Graphic

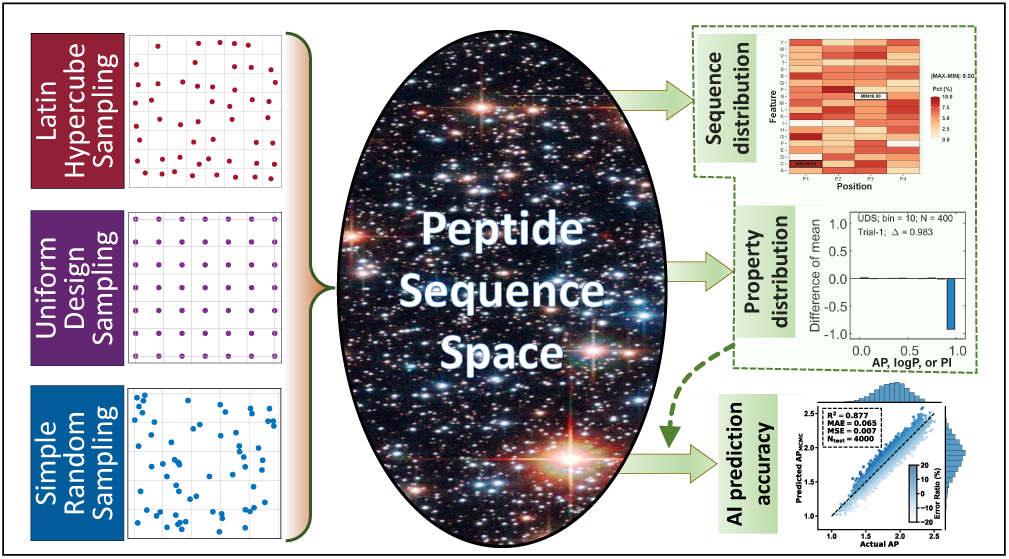

